# Mapping the Peaks: Fitness Landscapes of the Fittest and the Flattest

**DOI:** 10.1101/298125

**Authors:** Joshua Franklin, Thomas LaBar, Christoph Adami

**Author notes:** Co-First Authors (Joshua Franklin, Thomas LaBar).

## Abstract

**Background:** Populations exposed to a high mutation rate harbor abundant deleterious genetic variation, leading to depressed mean fitness. This reduction in mean fitness presents an opportunity for selection to restore adaptation through the evolution of mutational robustness. In extreme cases, selection for mutational robustness can lead to “flat” genotypes (with low fitness but high robustness) out-competing “fit” genotypes with high fitness but low robustness—a phenomenon known as “survival of the flattest”. While this effect was previously explored using the digital evolution system Avida, a complete analysis of the local fitness landscapes of “fit” and “flat” genotypes has been lacking, leading to uncertainty about the genetic basis of the survival of the flattest effect.

**Results:** Here, we repeated the survival of the flattest study and analyzed the mutational neighborhoods of fit and flat genotypes. We found that flat genotypes, compared to the fit genotypes, had a reduced likelihood of deleterious mutations as well as an increased likelihood of neutral and, surprisingly, of lethal mutations. This trend holds for mutants one to four substitutions away from the wild-type sequence. We also found that flat genotypes have, on average, no epistasis between mutations, while fit genotypes have, on average, positive epistasis.

**Conclusions:** Our results demonstrate that the genetic causes of mutational robustness on complex fitness landscapes are multifaceted. While the traditional idea of the survival of the flattest effect emphasized the evolution of increased neutrality, others have argued for increased mutational sensitivity in response to strong mutational loads. Our results show that both increased neutrality and increased lethality can lead to the evolution of mutational robustness. Furthermore, strong negative epistasis is not required for mutational sensitivity to lead to mutational robustness. Overall, these results suggest that mutational robustness is achieved by minimizing *heritable* deleterious variation.

## Background

All well-adapted populations receive an influx of deleterious variation every generation due to *de-novo* mutations. This influx reduces the population’s average fitness and causes a mutational load [2,11]. For populations with high mutation rates, deleterious variation can alter the dynamics of natural selection and lead to selection acting not just on individual genotypes, but on genotypes and their likely mutant genotypes [5,6,31,42]. This combination of a wild-type sequence and its mutants is known as a “quasispecies” and is thought to be a unit of selection in organisms such as the first RNA replicators [14, 15] and modern-day RNA viruses [27].

Since the quasispecies concept was first proposed [14,15], many studies have analyzed the conditions and consequences of evolutionary dynamics in this strong mutational regime (e.g., [4, 23, 42]). One consequence of evolution under high mutation rates is that populations are expected to evolve mutational robustness by fixing mutations that decrease the likelihood (and/or decrease the average effect) of deleterious mutations [12, 44]. In the extreme case, this trend can lead to genotypes with lower fitness out-competing genotypes with higher fitness if the lower-fitness genotypes are more mutationally-robust [37]. This effect has been termed the “survival of the flattest” effect, where “flat” genotypes, that is, those with lower fitness but greater mutational robustness, out-compete “fit” genotypes, meaning those with higher fitness but decreased mutational robustness [45]. While this effect was first observed in the digital evolution system Avida, it was later demonstrated with experimental evolution of RNA viruses [8,28,36].

Even though the evolutionary dynamics of the survival of the flattest effect are well-understood, the exact genetic causes of robustness are unknown. Indeed, the differences in the distribution of fitness effects [17] (that is, the local fitness landscapes) between fit genotypes and flat genotypes have not been explored. While it is generally assumed that mutational robustness evolves due to the evolution of an increased number of neutral mutational neighbors [18,39,41], other authors have argued that increased mutational sensitivity may evolve [3,23,26,32,46]. Additionally, recent work has indicated that populations evolving under strong genetic drift are expected to evolve “drift-robust” genetic architectures, giving rise to genotypes with a decreased likelihood of small-effect deleterious mutations and an increased likelihood of neutral and/or strongly-deleterious mutations [25]. A mapping of the local fitness landscapes of fit genotypes and flat genotypes is needed to compare the survival of the flattest effect to these other proposed evolutionary mechanisms.

To explore the local fitness landscapes of fit genotypes and flat genotypes, we first repeated the original survival of the flattest experiment. We then took genotypes adapted to either low or high mutation rates and obtained the distribution of fitness effects for these genotypes by examining mutants up to four substitutions away from the wild-type sequence. We found that flat genotypes had a greater likelihood of beneficial and neutral mutations (as expected for mutationally-robust genotypes) while fit genotypes had a greater likelihood of deleterious mutations, as expected from mutationally-sensitive genotypes. Contrary to expectations, we also found that flat genotypes had a greater likelihood of lethal mutations, suggesting that the survival of the flattest effect relies on a reduction of heritable deleterious mutation, which can be achieved by both an increase in neutral variations, as well as lethal mutations. These results illustrate the complexity of mutationally-robust genome architectures on non-trivial fitness landscapes.

## Materials and Methods

### Avida

Here, we describe the relevant details of Avida (version 2.14; our version, with added code for data analysis, is available at https://github.com/joshf93/MappingThePeaks) for the current experiments (see [33] for a full overview of the Avida software). In Avida, a finite population of computer programs (“avidians”) compete for the resources (memory, space, and processing time) required for reproduction. Each program consists of a circular genome of computer instructions that encode the ability to self-replicate. During this replication process, instructions may be copied inaccurately, resulting in mutations that are passed onto an avidian’s offspring. As avidians with faster replication speeds enter the population, they reproduce faster than slower-replicating avidians, leading to the spread of faster replicators and the extinction of slower replicators. In other words, there is selection for faster replicators. Therefore, because there is heritable variation and selection in Avida, populations of avidians undergo evolution by natural selection [34]. Avida has been used to test many concepts difficult to test in biological systems [1,10,19,20,24,29,38].

The Avida world consists of a toroidal grid of *N* cells; each cell can be occupied by at most one avidian and thus *N* is the maximum population size. During reproduction, offspring avidians are placed into one of the nine adjacent cells of its parent, including the cell that contains the parent. If some of these cells are unoccupied, a random empty cell is chosen. If all neighboring cells are occupied, a random cell is chosen, its occupant is removed, and the new avidian then occupies the cell. This random selection of replacement adds an element of genetic drift to Avida.

The limited number of cells in the Avida environment is one of the limiting resources over which avidians compete. The other limiting resource is the opportunity to execute the instructions in an avidian’s genome. The resource required to execute one instruction is called a “Single Instruction Processing” unit (SIP). During each unit of time in Avida (called an update) 30*N* SIPs are available to the population for execution. These SIPs are probabilistically distributed to avidians according to a figure of merit that is related to their ability to perform certain computations. These computations (binary logic operation) are the “traits” that avidians display. In a monoclonal population, all avidians express the same traits and thus have the same merit and thus receive on average 30 SIPs (i.e., they execute 30 genome instructions during that update). However, avidian lineages can evolve the ability to increase merit, and thus increase the number of instructions that can be executed per update.

Fitness in Avida is implicit and can only be estimated by executing the computer instructions within an avidian’s genome. In this regard, Avida differs from classical evolutionary simulations where genotypes are assigned a fitness and their frequency increases or decreases proportionally to this assigned value. Here, faster replicators out-compete slower replicators and fitness emerges from this dynamic. There are two means by which an avidian lineage can evolved increased replication speed. First, genotypes that require fewer instruction executions to undergo reproduction will reproduce faster than avidians that require a greater number of instruction executions. In environments where genome size can vary, this is often achieved by decreasing genome size. In fixed genome size environments, this trajectory of fitness gain still occurs, often through the fixation of additional instructions that copy genome information from parent to offspring. The second methods of fitness increase in Avida is through the evolution of complex adaptations: the ability to perform one- and two-input Boolean logic calculations (the traits discussed earlier). In the setup used here, nine of these functions are potentially rewarded with SIPs; the amount of SIPs awarded is proportional to the complexity of the evolved trait [30]. The performance of these calculations increases an avidian’s merit. In other words, the more calculations an avidian can perform, the more SIPs it receives per update and it can then execute a larger portion of its genome per unit time. This increases the avidian’s replication speed and leads to selection for the ability to perform these calculations. When Avida estimates a genotype’s fitness, it is estimated as the genotype’s merit divided by the number of instruction executions required for replication (otherwise known as its gestation time).

Avidian populations evolve these complex adaptations by fixing a sequence of mutations. This genetic variation is introduced into a population when an avidian copies one instruction inaccurately into its daughter’s genome. The rate at which these mutations occur is set by the experimenter and is usually set as a rate “per instruction copied” (a copy-mutation rate). When mutations occur during this copying process, multiple mutations can occur per replication event. Additionally, in experiments where genome size can evolve, insertion and deletion mutations occur upon division (not during the replication process) at a pre-specified rate. These indel mutations insert or delete one instruction. Finally, large-scale genome changes can occur, but these changes are caused by the specific replication algorithm encoded by an avidian’s genome; these mutations are inherent to the Avida genetic code (implicit mutations) and are not under the control of the experimenter.

### Experimental Design

Here, we repeated the experimental protocol of the original survival of the flattest study [45]. We differed from the original study in one aspect: the placement of offspring. Here, we used Avida’s current default setting, where a new offspring randomly replaces one of the nine neighbors of its parent (including possibly the parent). The original paper used mass-action reproduction, where new offspring could randomly replace any offspring in the population. We first generated a collection of genotypes during the *Initial Adaptation* step. We evolved 201 populations of 3,600 individuals for 5 10^4^ updates. All populations initially started with the default Avida ancestor with an one-hundred instruction genome. Point mutations occurred at a rate of 0.0075 mutations per instruction copied and insertion/deletion mutations occurred at a rate of 0.05 mutations per division each. At the end of these populations’ evolution, we extracted the most abundant genotype.

For the *Mutation Rate Adaptation* phase of the experiment, we duplicated each of the 201 genomes and used one as progenitor of an experiment at a fixed *low* genomic mutation rate of 0.5 mutations per genome per generation, and one at a fixed high genomic rate of 2.0 mutations genome per generation. To achieve this, we adjusted the per-site mutation rate of each genome so that the targeted genomic rate was achieved, and disallowed sequence length changes. In this manner, the *Initial Adaptation* created genetic variation used as input to the adaptation step at a fixed genomic rate. From this stage onward each genotype was limited to the tasks it could complete at the end of the *Initial Adaptation* phase, i.e., they were not allowed to evolve novel traits, though they could lose traits they already possessed. Each of the populations adapting to high or low mutation rate consisted of 3600 individuals and evolved for 10^4^ generations. After evolution we again selected the most abundant genotype from each population and denoted genotypes from the low mutation environment “fit” genotypes, and genotypes from the high mutation environment “flat” genotypes.

Next, we performed competition experiments between each pair of fit and flat genotypes (each pair has the same ancestor). We only performed these competitions with genotype pairs where the fit genotype was 1.5 times more fit than the flat genotype. For the competitions, we seeded populations of 3600 individuals with 1800 individuals of each genotype and ran the simulation for 200 generations. We repeated these competitions across a range of genomic mutation rates (0.5 to 3.0 mutations/genome/generation in increments of 0.5) and performed five replicates per mutation rate. We tracked the abundance of the fit genotypes over time to determine the outcome of the competition. We classified the flat genotype as the winner if it reached at least 95% of the population in three of five replicates at a given mutation rate.

### Data Analysis

We performed mutational analyses to determine the fitness landscapes of the fit and flat genotypes used in the competitions using Avida’s “analyze” mode. In analyze mode, avidians run through their life-cycle in isolation (as opposed to in a population) and characteristics such as their fitness can be estimated. We first estimated the base fitness of all avidian genotypes used in the experiment. Then, we took the genotypes used in the competition and generated all possible single mutants. For each genotype, there are 25*L* single mutants, where *L* is the genome size and 25 is the number of possible alternative alleles at each position in the genome, as there are 26 instructions in the Avida instruction set. We then repeated this procedure for all double mutants. For triple- and quadruple-mutants, we randomly sampled 10^8^ mutants as the number of all possible mutants made generating every mutant computationally prohibitive^1^.

After generating these mutants, we calculated their fitness and compared them to their ancestor genotype. Mutants with greater fitness were designated as beneficial and mutants with the same fitness were designated as neutral. Mutants that were viable but with decreased fitness were designated as deleterious. Mutations resulting in genomes that could not reproduce were designated as lethal. To estimate differences in epistasis between fit and flat genotypes, we fit the mutational analysis data to the following equation:

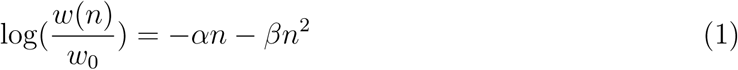

Here, *n* is the genetic distance from the initial genotype, *w*(*n*) is the mean fitness of all mutants at a given genetic distance, and *w*_0_ is the fitness of the initial genotype. This is a similar equation to that fit in the original survival of the flattest paper [45], with the mean fitness estimated at a given mutation rate substituted by the mean fitness of all n-mutants (as is often used [29]). *α* and *β* can be thought of as parameters that measure a genotype’s mutational robustness and epistasis, respectively. A greater *α* indicates that a genotype is less mutationally-robust, while a greater *β* indicates epistasis is more negative (i.e., the fitness effect of a double-mutant is less than expected from the fitnesses of the two single-mutants).

All statistical analyses were performed using R 3.4.3 [35] and figures were generated with either the ggplot2 R package [40] or the Matplotlib Python package [21].

## Results

We first evolved 201 populations and isolated the most abundant genotype from each population. We then evolved these genotypes in both a high mutation rate environment and a low mutation rate environment and isolated the most abundant genotype from each environment. The genotypes from the low mutation rate environment did not significantly change in fitness compared to their ancestor (Fig. 1a; Wilcoxon Rank Sum test, *p* = 0.411). The genotypes from the high mutation rate environment significantly decreased in fitness compared to their ancestors (Fig. 1a; *p* = 2.216×10^*−*7^). By tracking the fitness of the high-mutation-rate populations through time, we observe that these genotypes first severely decrease in fitness upon transfer, but then recover fitness as they search out new, and presumably more mutationally robust, fitness peaks (Fig. 1b).

**Figure. 1:**
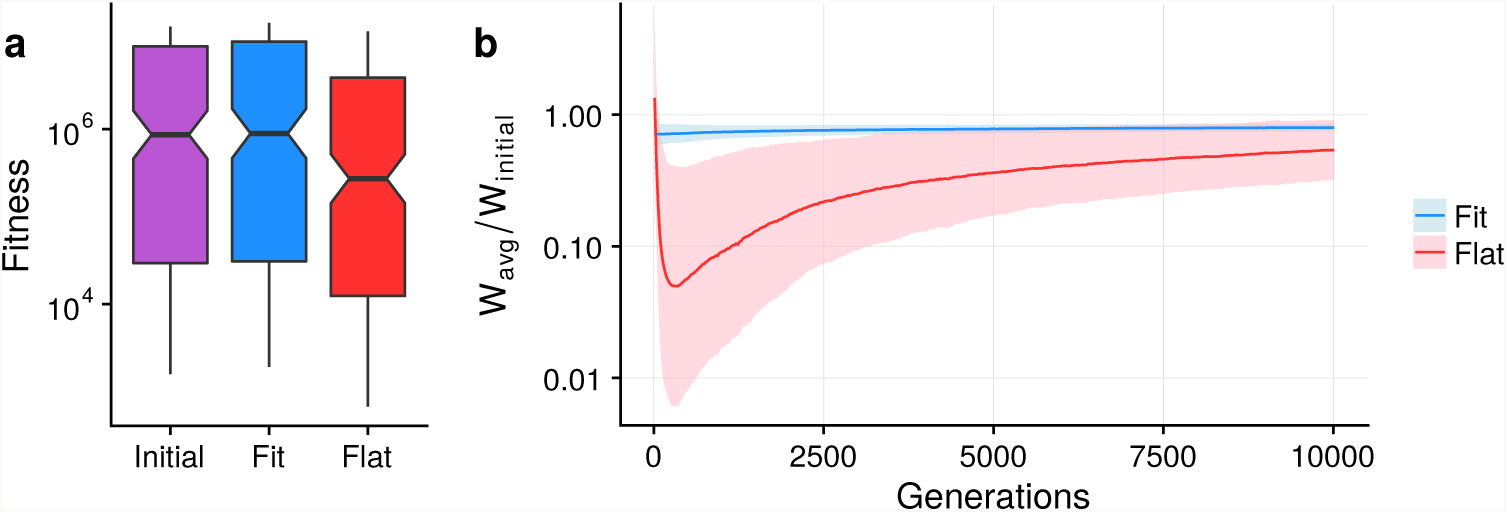
Populations adapt to different fitness peaks in high mutation rate environments. a) Fitness values for the most abundant genotype from each population after the *Initial Adaptation* experiment (denoted by “Initial”) and after adaptation to low (“Fit”) and high (“Flat”) mutation rate environments. For boxplots, lines are the median values, boxes show the interquartile range, and whiskers are 1.5 times the interquartile range or the minimum/maximum value; points are outliers. Overlapping notches suggest that the difference in medians is not significant. b) Relative average fitness values over the course of the *Mutation Rate Adaptation* experiment. Lines are the mean value across all populations and the shaded regions are standard deviations. Blue represents fit genotypes and red represents flat genotypes.

Of our 201 pairs of genotypes adapted to low mutation rates and high mutation rates, 166 pairs evolved such that the low mutation rate genotype (the fit genotype) had a fitness greater than 1.5 times the high mutation rate genotype (the flat genotype), which is the threshold used in the original study [45]. We took these pairs and performed competition experiments between the two genotypes across a range of mutation rates. In 76 (45.8%) of these competition experiments we noted that the eventual winner of the competition depended on the mutation rate, and that a switch between winners occurred at a critical mutation rate, just as was observed in the initial study [45]. Fit genotypes win the competition at low mutation rates, but flat genotypes win the competition at high mutation rates even with the large fitness difference (Fig. 2). It is worth noting that our definition of winning excluded many genotype pairs that showed the effect to a lesser degree.

**Figure. 2:**
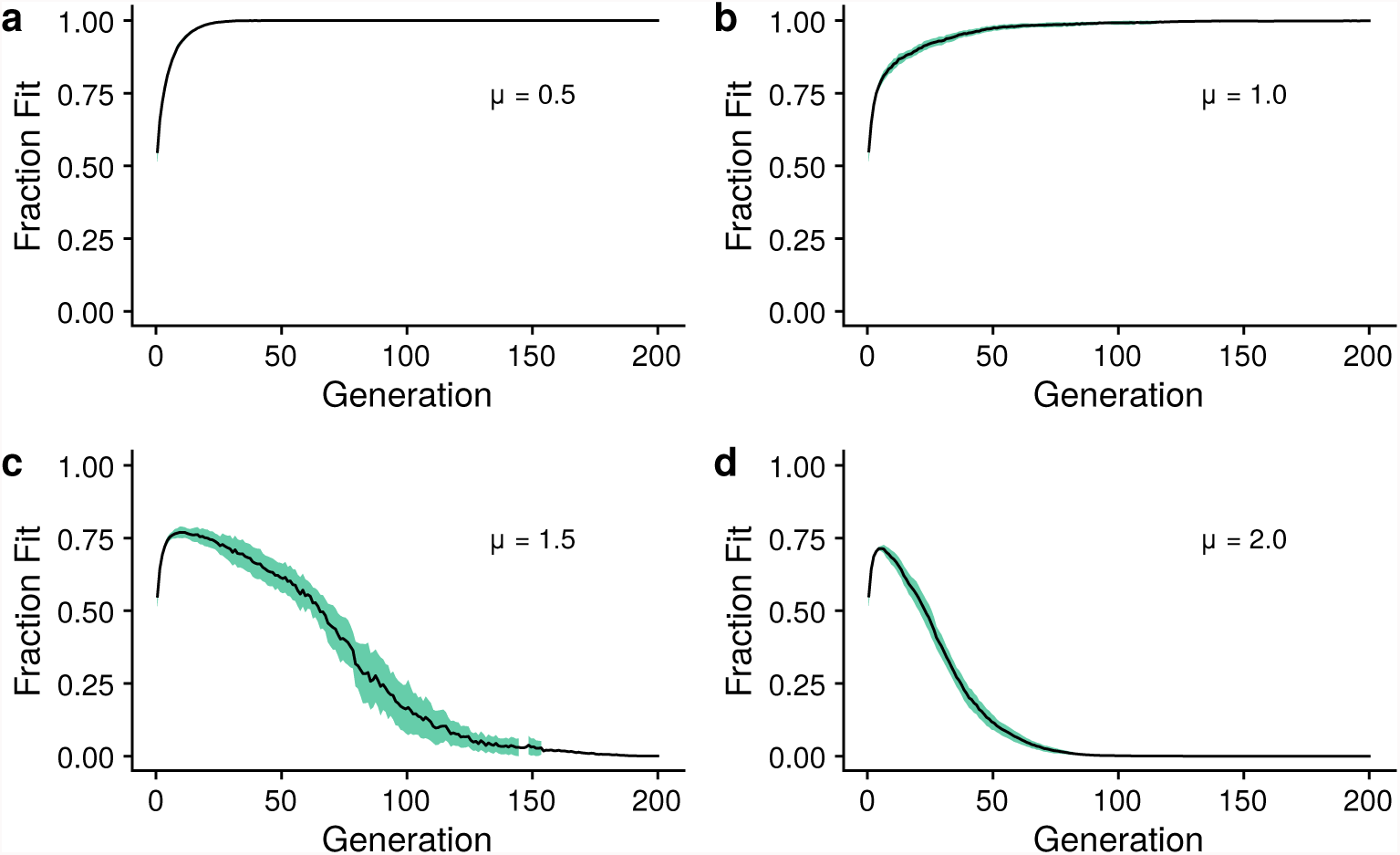
Competition outcome between fit and flat genotypes is determined by the mutation rate. Outcome from competition experiments between one fit/flat genotype pair. Black line is the mean fraction of the population consisting of the fit genotype as a function of time (measured in generations). Green shading represents one standard deviation. Each subplot shows the results for a given genomic mutation rate for five replicates.

Next, we analyzed the mutational neighborhood (the local fitness landscape), of all 166 pairs of fit and flat genotypes. We first generated all genotypes containing one point mutation and measured their fitness. There are significant differences in particular classes of mutations. Flat genotypes have a greater likelihood of beneficial (Fig. 3a; Wilcoxon Rank Sum test, *p* = 1.35 × 10^−20^), neutral (Fig. 3b; *p* = 2.66 × 10^−35^), and lethal (Fig. 3d; *p* = 0.0168) mutations. Fit organisms have a greater likelihood of deleterious mutations (Fig. 3c; *p* = 1.31 × 10^*−*44^). Additionally, fit genotypes have lower mean relative fitness than flat genotypes (Fig 3d; *p* = 2.24 × 10^*−*27^). We also analyzed the mutational neighborhood for sequences that are two, three, or four mutations away from the wild-type sequence and found similar results to the one-mutant mutational neighborhood (Fig. 4).

**Figure. 3:**
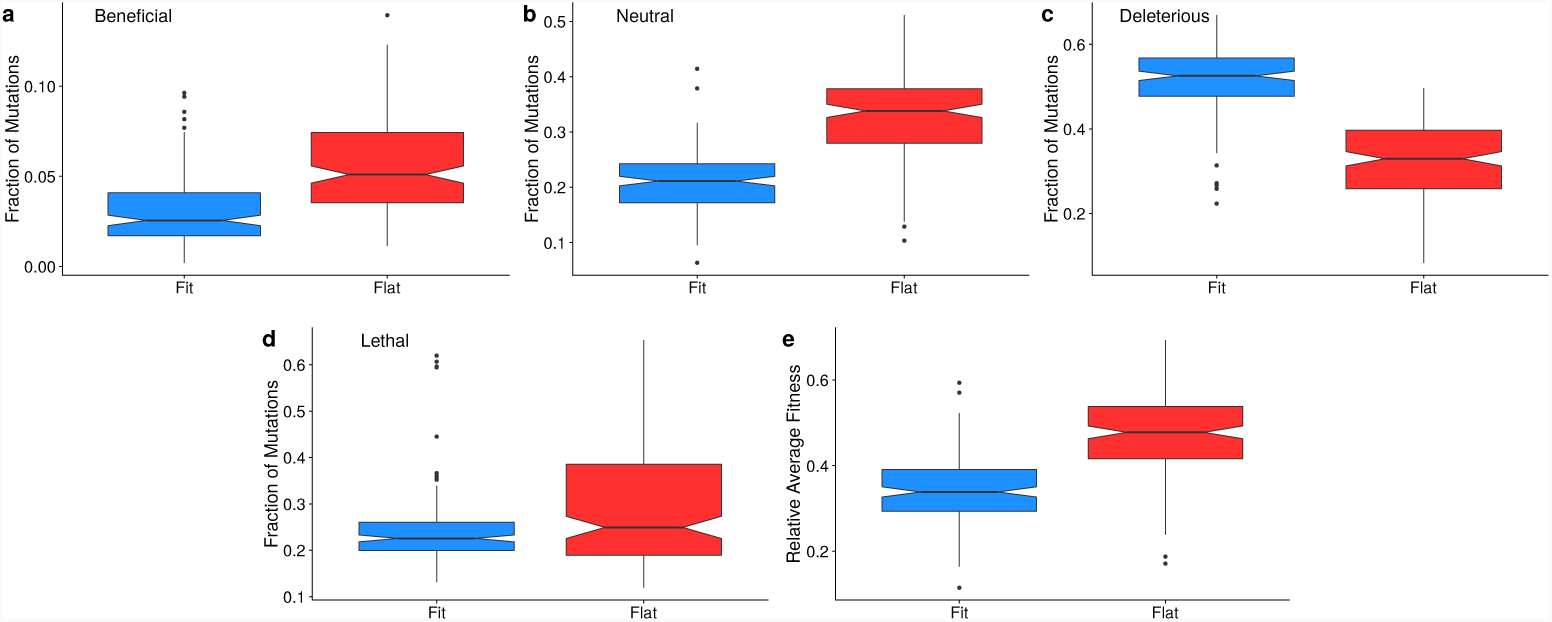
Description of local fitness landscapes for fit and flat genotypes. a-e) Distribution of fitness effects with mutations grouped by their broad effect. Boxplots as previously described. Lineage refers to whether the data is for fit genotypes (blue) or flat genotypes (red). f) Average relative fitness of all one-step mutants for each fit and flat genotype.

**Figure. 4:**
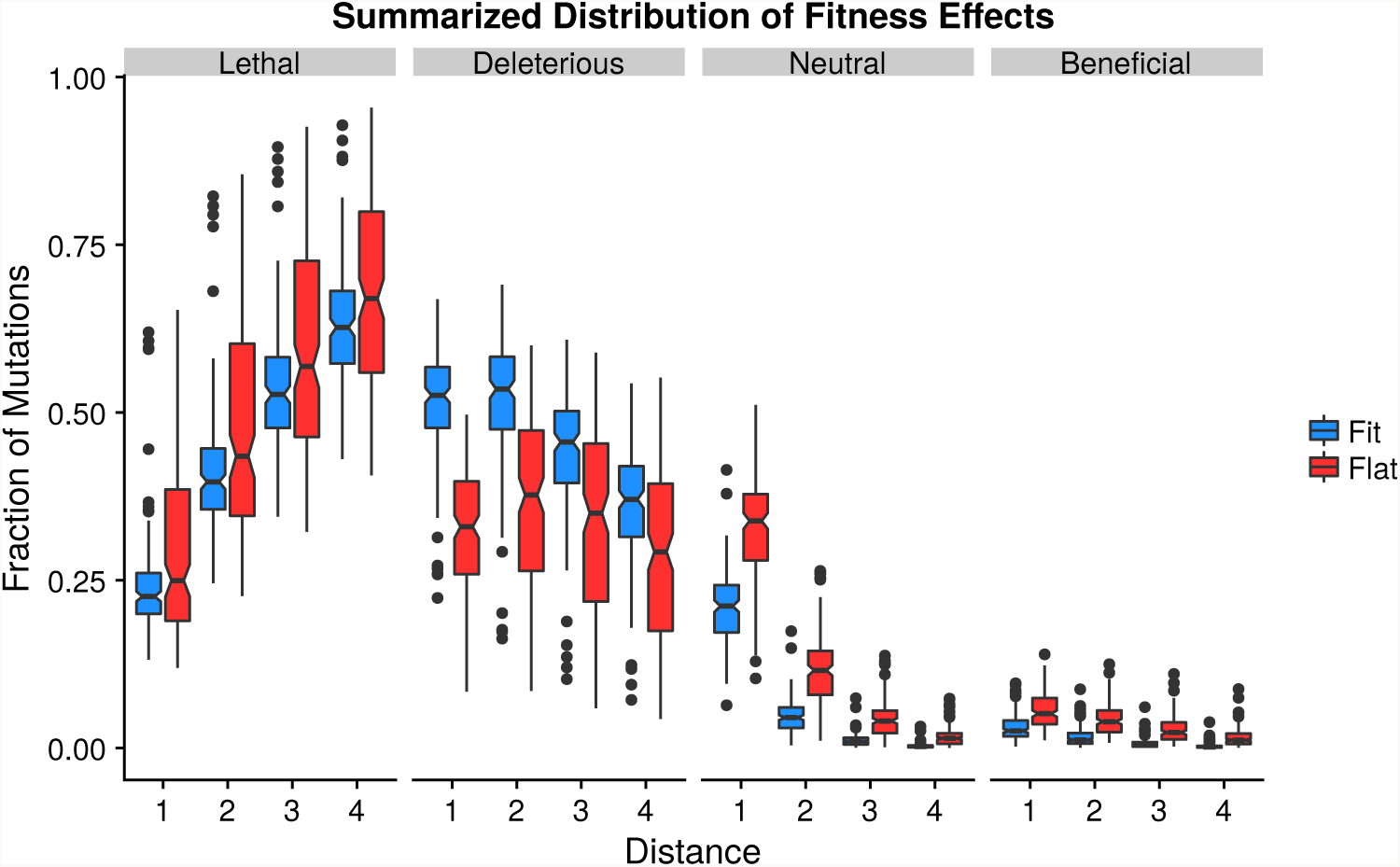
Summarized distribution of fitness effects for mutants with multiple mutations. Proportion of mutations with a mutational effect classification for mutants up to four mutations away from the wild-type sequence. See text for definitions of different classes of mutations. Distance is the number of mutations away from the wild-type sequence. Data for mutants with distance of one is the same as in Figure 3. Blue (red) boxplots are the proportions from the fit (flat) genotypes. Boxplots as previously described.

Finally, we used the mutational neighborhood data for sequences up to four mutations away from the wild-type sequence to estimate differences in epistasis between fit and flat genotypes. We fit the relative average fitness values as a function of mutational distance to a curve containing two parameters (see Methods for further details): *α* (as a measure of mutational robustness) and *β* (as a measure of epistasis). As expected from the data on the relative fitness of one-step mutants (Fig. 3e), fit genotypes have greater alpha values (Fig. 5a; *p* = 6.88 × 10^*−*28^). Flat genotypes have greater *β* values (Fig. 5b; *p* = 1.84 × 10^−34^), indicating that flat genotypes have greater negative epistasis than fit genotypes. However, it should be noted that flat genotypes have, on average, almost no epistasis, while fit genotypes have positive epistasis. These data demonstrate how the selection pressure for mutational robustness drives populations to alternative areas of the fitness landscape.

**Figure. 5:**
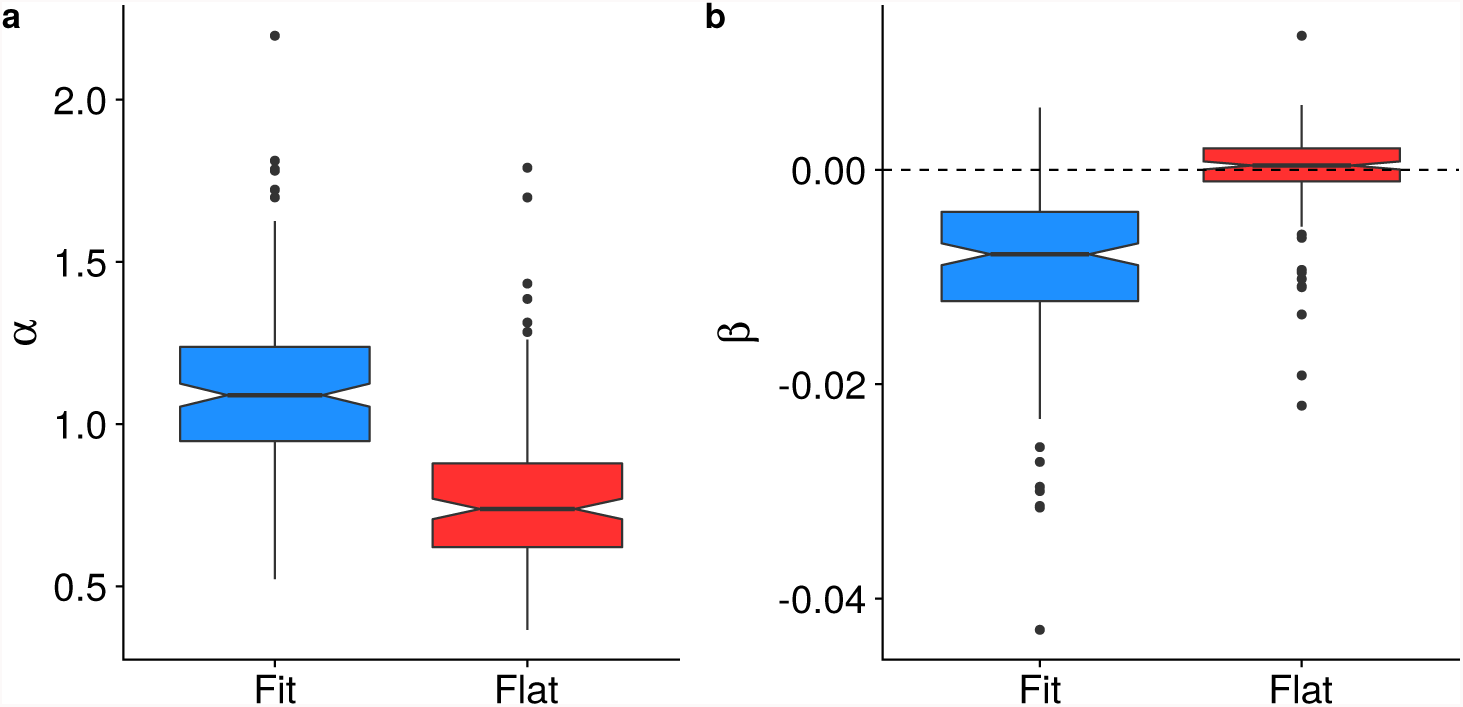
Distribution of *α* and *β* values for fit and flat genotypes. Distributions of *α* and *β* for each treatment. *α* is a measure of single-mutant fitness, while *β* is a measure of epistasis (see Methods for further details). Boxplots as previously described.

## Discussion

We used digital experimental evolution to map the local fitness landscape of high fitness, but mutationally-fragile,“fit” genotypes adapted under low mutation rates, and low fitness, but mutationally-robust, “flat” genotypes adapted under high mutation rates. After repeating the original survival of the flattest effect [45], we calculated the distribution of fitness effects for *de-novo* point mutations up to four substitutions from the wild-type genotypes. We found that, as expected, flat genotypes were more mutationally robust than fit genotypes. Flat genotypes had a lower likelihood of deleterious mutations and a greater likelihood of neutral and beneficial mutations than fit genotypes. Surprisingly, flat genotypes also had a greater likelihood of lethal mutations and more negative epistasis than fit genotypes, suggesting that the factors responsible for mutational robustness involve only limiting *heritable* deleterious variation.

Around the time when the original survival of the flattest paper first appeared, mutational robustness was largely conceptualized as a decrease in the likelihood of a deleterious mutation [39, 45]. And while these changes in genetic architecture can lead to mutational robustness, work since that original publication has suggested that *increases* in the effect of deleterious mutations, not decreases, can also lead to robustness [3,32]. Increased severity of deleterious mutations can lead to population-level robustness as purifying selection is more effective against strongly-deleterious mutations. It has also been proposed that increased negative (or synergistic) epistasis can evolve as a mechanism to increase the strength of purifying selection and hence robustness [26, 46].

Our results suggest that all of these genetic mechanisms (increased neutrality, increased severity of deleterious mutations, and increased negative epistasis) can contribute to mutational robustness. These data support a previously-proposed relationship between increased neutrality (low *α*) and increased negative epistasis (high *β*), based on data from Avida, RNA-folding, and neutral network fitness landscapes [43]. In other words, a population can only maximize immediate mutational robustness at the cost of increased deleterious severity of mutants farther away in the fitness landscape. We should also note that mutations on a “flat” genetic background are not, on average, negatively epistatic. Flat genotypes evolve to have, on average, no epistasis between mutations, as previously shown to occur in high mutation rate Avida populations [13]. However, they are more negatively-epistatic than fit genotypes, which have positive epistasis between mutations. The lack of negative epistatic interactions in flat genotypes is likely related to their abundance of lethal mutations. Additional mutations on a genetic background with a lethal mutations cannot, by definition, lead to a greater decline in fitness (negative epistasis).

Theoretical arguments suggest that increasing the severity of mutations not only protects against high mutation rates, but also can protect against fitness loss due to genetic drift [22], although this depends on the specific distribution of fitness effects [7]. Recently, LaBar and Adami proposed the concept of “drift robustness,” or the idea that small populations tend to populate drift-robust fitness peaks with a low likelihood of slightly-deleterious mutations (and a high likelihood of both neutral and strongly-deleterious mutations), while large populations would adapt to drift-fragile fitness peaks with a high likelihood of slightly-deleterious mutations [25]. Both flat genotypes and drift-robust genotypes have some noticeable similarities in their distribution of fitness effects (increased neutrality, in particular), so it is worth asking whether the evolution of drift robustness and survival of the flattest is really the same phenomenon.

The main difference between drift robustness and survival of the flattest is the role of natural selection in the two phenomena. The evolution of drift robustness does not require competition between drift-robust and drift-fragile genotypes [25]. Instead, small populations tend to evolve towards drift robustness because they can only maintain fitness on drift-robust genetic backgrounds. Survival of the flattest, however, occurs when a flat genotype out-competes a fit genotype, and is driven by selection for mutational robustness. It has also been established that the critical high mutation rate above which this selective advantage occurs is independent of population size [9], while drift robustness, almost by definition, is strongly dependent on population size. Furthermore, our previous work demonstrated that drift-robust genotypes are not more mutationally-robust than drift-fragile genotypes, if one measures mutational robustness as the mean relative fitness of all single mutants [25]. Single mutants of flat genotypes are more fit, on average, than those of fit genotypes, again illustrating a difference between drift robustness and survival of the flattest. Thus, while similarities exist between drift-robust genetic architecture and “flat” genetic architecture, it appears that they represent distinct evolutionary responses to different population-genetic environments.

It would be worthwhile to determine if the results observed held here can be replicated with RNA viruses, the biological model for the survival of the flattest effect [8,28,36]. It would be illuminating to test if the likelihood of lethal mutations does increase in these evolved viruses, especially in light of studies arguing that RNA viruses have high mutational *fragility*, not high mutational robustness [16]. Such a study would lead to a better understanding of the mutational mechanisms that lead to robust organisms in nature.

## Conclusions

In high mutation rate environments, populations will evolve alternative genetic architectures that increase their robustness to deleterious mutations. Classically, this is viewed as a survival of the flattest effect [45], where populations adapt to fitness peaks with increased neutrality [39]. Here, we repeated the original experiments to finely map the fitness landscapes of fit and flat genotypes. Flat, mutationally-robust genotypes adapt to fitness peaks with an increased likelihood of both neutral genotypes and lethal genotypes, a decreased likelihood of deleterious mutations, and greater negative epistasis compared to fit genotypes. These results demonstrate the genetic complexity of mutationally robustness genomes that minimize the impact of heritable deleterious genetic variation.

## Ethics approval and consent to participate

Not applicable.

## Consent for publication

Not applicable.

## Availability of data and material

All Avida configuration files and data analysis scripts to recreate these experiments, analyze the data, and recreate the figures are available at https://github.com/joshf93/MappingThePeaks.

## Author’s contributions

JF performed the experiments and analyzed the data. JF, TL, and CA designed the study and wrote the manuscript.

## Acknowledgements

T.L. acknowledges a Michigan State University Distinguished Fellowship, a BEACON fellowship, and the Russell B. DuVall award for support. This work was supported in part by Michigan State University through computational resources provided by the Institute for Cyber-Enabled Research.

## Footnotes

1 The number of possible *n*-mutants of a length *L* genome encoded in an alphabet of size 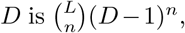, which translates to over 1.5 *×* 1012 possible mutants for *L* = 100, *n* = 4, and *D* = 26.

